# Long-term homeostasis in microbial consortia via auxotrophic cross-feeding

**DOI:** 10.1101/2025.01.08.631749

**Authors:** Nicolas E. Grandel, Amanda M. Alexander, Xiao Peng, Caroline Palamountain, Razan N. Alnahhas, Andrew J. Hirning, Krešimir Josić, Matthew R. Bennett

## Abstract

Synthetic microbial consortia are collections of multiple strains or species of engineered organisms living in a shared ecosystem. Because they can separate metabolic tasks among different strains, synthetic microbial consortia have myriad applications in developing biomaterials, biomanufacturing, and biotherapeutics. However, synthetic consortia often require burdensome control mechanisms to ensure that the members of the community remain at the correct proportions. This is especially true in continuous culture systems in which slight differences in growth rates can lead to extinctions. Here, we present a simple method for controlling consortia proportions using cross-feeding in continuous auxotrophic co-culture. We use mutually auxotrophic *E. coli* with different essential gene deletions and regulate the growth rates of members of the consortium via cross-feeding of the missing nutrients in each strain. We demonstrate precise regulation of the co-culture steady-state ratio by exogenous addition of the missing nutrients. We also model the co-culture’s behavior using a system of ordinary differential equations that enable us to predict its response to changes in nutrient concentrations. Our work provides a powerful tool for consortia proportion control with minimal metabolic costs to the constituent strains.

## Introduction

As the complexity of synthetic biology constructs continues to increase, researchers are moving away from single-strain systems (1). Multi-strain microbial consortia enable division of labor, which reduces the metabolic load on individual organisms and allows for the entire consortium to behave more efficiently (2–4). This benefit has led to advances in microbiome engineering (5–8), bioremediation (9–14), and bioproduction (2, 3, 15–18), making consortia engineering one of the fastest growing fields in synthetic biology. Most natural microbial communities are made of many species working together. These consortia exhibit complex collective behaviors such as patterning (19–21), increased robustness (22– 25), and co-evolution through horizontal gene transfer (26– 28). While there are benefits to engineering natural communities, the complexity of interactions within these systems often makes this infeasible. Synthetic consortia offer the advantage of being simple while retaining the benefits of the division of labor found in natural communities. This makes synthetic consortia ideal both for increasing the efficiency of known biological solutions and for studying behaviors only seen in complex communities. The chief challenge, then, becomes creating those same emergent behaviors through rational design.

One of the largest barriers to synthetic consortia design is the problem of competitive exclusion. Without pressure to coexist, simple differences in growth rates lead to the loss of less fit consortium members (29). Even worse, many desirable behaviors (e.g., bioproduction (4, 16, 17)) rely not just on coexistence but on specific population ratios. Maintaining such specific ratios requires precise control of each strain’s growth rate. Further complicating the matter, any mutation that increases fitness will break this control, making longterm ratio control difficult. An ideal ratio control strategy would be robust over time and suitable for a wide variety of applications.

Many existing population control strategies maintain member coexistence but lack in other design pillars. Broadly, these strategies define pairwise interactions to control the growth rate of all members (1, 30). Toxin-antitoxin systems and antimicrobials have proven promising in a wide variety of contexts, but are sensitive to mutation and rely on the production of molecules that limit growth (31, 32). Symbiotic relationships are simple, but are thought to be limited by the necessity of strict inter-dependence (33–35). Quorum sensing regulation has been popular for enabling tunability in recent years, but the limited number of existing quorum sensors in turn limits the number of applications that this method can be combined with (36–39). Mutualistic auxotrophy was one of the first population control methods used in synthetic microbial consortia (33). While it has been known for some time that this commensalism is capable of maintaining multiple members of a population in a variety of contexts (34, 35, 40), little has been done to study the stability or tunability of this control strategy due to the slow growth rates of auxotrophs.

Here, we present a simple and tunable mechanism for controlling consortia ratios using mutualistic *E. coli* auxotrophs. The ratio is stabilized by the excess production of the cross-fed metabolites and is tunable via the exogenous addition of each metabolite. We show that auxotrophs in the consortium converge to a stable population ratio in continuous co-culture as a result of balanced growth rates, and that this ratio is insensitive to inoculation conditions. We further show that we can tune this ratio by supplementing the deficient amino acids and increasing the growth rate of a constituent strain. We develop and validate a mathematical model that explains the mechanisms of co-culture dynamics and ratio control. When fit to data this model predicts coculture ratios under novel conditions. Our results represent a powerful new tool for controlling microbial community composition.

## Results

### System description

To test whether cross-feeding auxotrophs can regulate their relative population abundance in co-culture (Figure 1*A*), we co-cultured two strains taken from the Keio collection (41), Δ*argC* and Δ*metA*. These strains have a kanamycin resistance marker in the place of the *argC* or *metA* genes, which are required for arginine or methionine production, respectively. They have been previously shown to grow mutualistically in minimal media via cross-feeding (33). To do so, the Δ*argC* strain produces excess methionine, which allows the Δ*metA* strain to grow and, in turn, produce excess arginine, allowing the Δ*argC* strain to grow.

**Fig. 1.**
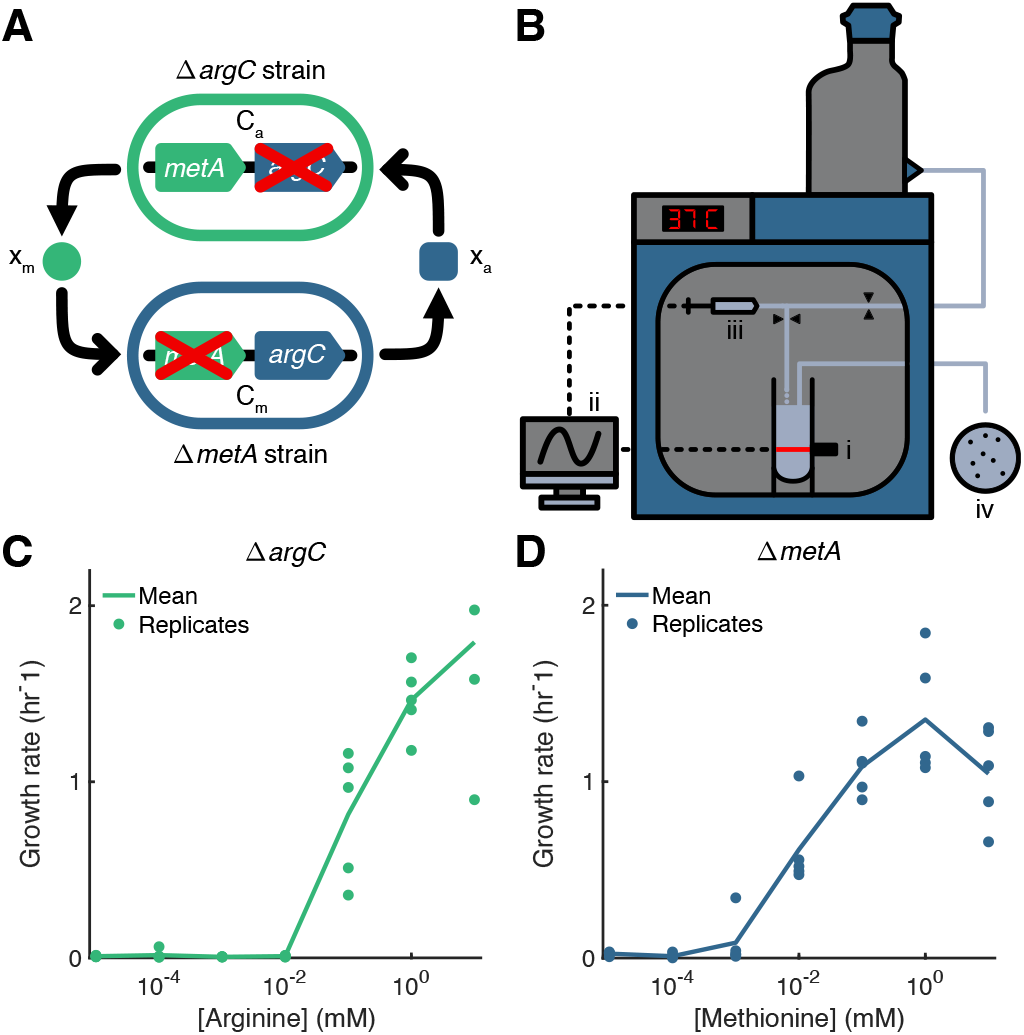
A simple, tunable system for ratiometric control using mutualistic auxotrophs. **(A)** Schematic of control system and corresponding model state variables. Two strains of *E. coli* containing chromosomal deletions in either the *metA* (*C*_*m*_) or *argC* (*C*_*a*_) genes are grown together in minimal media, where growth is only possible by the mutualistic cross-feeding of methionine (*x*_*m*_) and arginine (*x*_*a*_). **(B)** Schematic of continuous culture turbidostat setup. **(i)** OD readings of the cell culture are taken periodically. **(ii)** OD data are used to compute necessary dilution volumes to maintain a set OD at each time step. **(iii)** A mechanized syringe and pinch valve system pulls the calculated dilution volume of media and dispenses it into the culture tube. **(iv)** Every 6 hours, dilution effluent is plated onto selective and nonselective media plates to estimate the population ratio. **(C)** and **(D)** Bulk culture growth rates of the Δ*argC* and Δ*metA* strains as arginine or methionine is added to minimal M9 media, respectively. Each strain’s independent growth rate is tunable via the addition of its corresponding missing metabolite.

In any microbial consortium, the abundance of each constituent strain is dependent on its growth rate. Growth regulation is thus essential to ratio control, but difficult in practice due to its sensitivity to mutation. Most ratio control mechanisms rely on limiting each member strain’s growth, making any random increase in fitness fatal to the precise control of the entire system. We noted that the mutualistic relationship between the Δ*argC* and Δ*metA* strains could overcome this sensitivity because of its robustness to potential mutations. Since each strain contained a whole-gene chromosomal deletion, it is unlikely that either strain would regain its metabolic function. However, ratio control also requires the ability to tune the growth rate of each strain to modulate their abundances. Beyond the cross-feeding capability, both strains exhibit robust growth in co-cultures, indicating that their growth rates in this environment are not substantially different from each other (33). Such similar growth rates indicate that, if their growth rates could be tuned, it should be possible to create a wide range of relative abundances. We hypothesized that the growth rate of each strain in this mutualistic system was determined by the availability of either arginine for the Δ*argC* strain or methionine for the Δ*metA* strain.

To test if we could control the strain growth rates to tune their relative abundances using these nutrients, we grew each strain in minimal media monoculture at a range of either arginine or methionine concentrations varying from 10 nM to 10 mM (Figure 1*C* and *D*). After overnight growth, each strain was inoculated into supplemented M9 media at equivalent cell densities as measured by OD_600_. We monitored the cell density of each culture via plate reader and used the changes in OD over time to estimate the growth rate under each growth condition. In both cases, the strain’s growth rate was restored to doubling times of under one hour at the highest concentrations, demonstrating the dominant effect of the missing metabolite on each auxotroph’s growth rate. Additionally, each strain exhibited a wide range of growth rates, indicating the possibility for a wide range of co-cultured ratios.

### System reaches steady state

With confirmation that the growth rates of these two strains could be tuned by the presence of either arginine or methionine, respectively, we next tested their ability to regulate their relative abundance in coculture. To do so, we co-cultured the Δ*argC* and Δ*metA* strains in a continuous culture turbidostat designed to maintain a constant OD_600_ over the course of the experiment (Figure 1*B*) (42). By periodically recording the culture OD and comparing it to an established setpoint, the turbidostat could add reserved minimal M9 media to maintain the setpoint OD as necessary. Culture media in excess of 15 mL was allowed to flow into a waste container as effluent. We periodically collected and plated the individual turbidostat dilutions on plates containing either rich media or minimal media supplemented with methionine to estimate the relative abundance of the Δ*metA* strain over the course of several days. We also kept a running log of both OD measurements and dilution volumes to estimate the co-culture growth rate.

To test whether the consortium converged to a stable ratio, we inoculated the strains at a range of initial ratios and monitored their relative abundances in continuous culture. The consortium came to around a ratio of 3:1 (Δ*metA*: Δ*argC*) within 24 hours (Figure 2*A-C*). This ratio persisted over several days after stabilizing, and the community gave no indication of losing fitness (Figure 5*A-C*). We found that 99:1 and 1:99 OD ratio inoculations did not alter either the final ratio or the time to steady state (Figure 2*B* and *C*). This indicates that this ratio control mechanism is robust and does not require precise estimation of relative strain abundances prior to co-culturing.

**Fig. 2.**
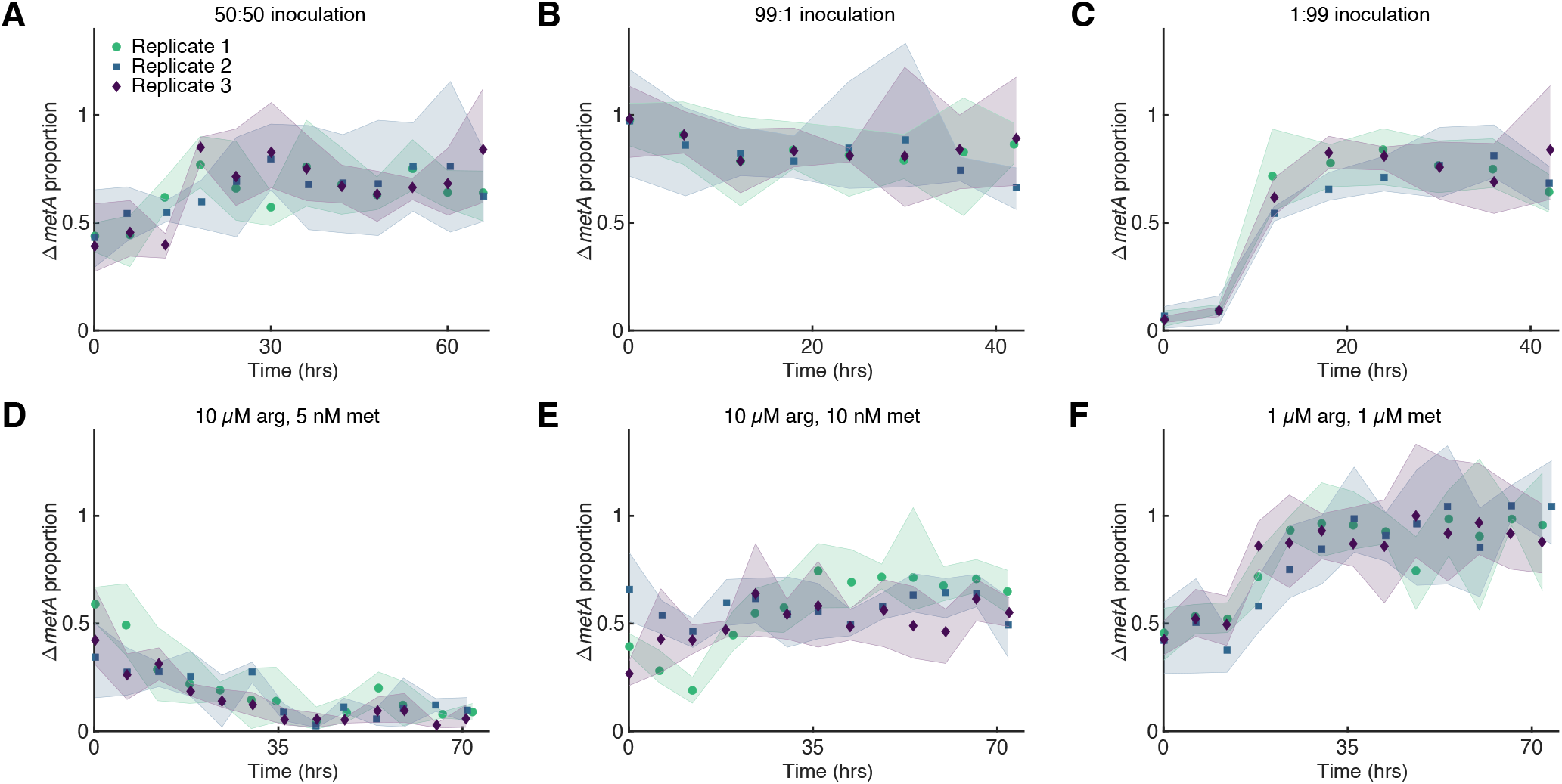
Mutualistic auxotrophy creates a stable population ratio in minimal media. **(A)** Δ*metA* population proportion in continuous culture of minimal M9 media. Cultures inoculated at ∼50:50 OD initial ratio. **(B)** and **(C)** Δ*metA* population proportion in continuous culture of minimal M9 media. Cultures were inoculated at either 1:99 or 99:1 OD to test robustness of steady state stability. **(D-F)** Tuning the Δ*metA* population proportion in continuous culture with the addition of arginine and methionine. We added both methionine and arginine to the turbidostat setup, allowing us to tune the strain ratio above and below our initial unsupplemented experiments. Proportions calculated as ratio of triplicate colony forming units counted on selective LB plates (total population) and M9 100 *µ*M methionine plates (Δ*metA* population). Icons represent ratio of mean colony counts, shaded areas represent maximum and minimum of nine possible combinations of two sets of triplicates. Ratios of *>* 1 are possible due to the possibility of growing more colonies on the M9 plate than on the LB plate.

### System tunability

We next attempted to tune the steady state ratio of the system by supplementing the minimal media with arginine and methionine (Figure 2*D-F*). Cells were grown under identical conditions to the prior experiments, except for added arginine or methionine at indicated amounts. While neither strain needed the other for its survival, as long as each experienced a sufficient growth benefit from the presence of the other, the co-culture could still reach a steady state that maintained both strains. This method allowed us to produce a wide range of stable population ratios. These included concentrations which allowed either strain to make up the majority (∼90% of the total population) (Figure 2*D* and *F*). Finer grained changes were also possible by adjusting the amount of added arginine and methionine. For instance, we were able to lower the ratio from the baseline ∼75% Δ*metA* to ∼50% Δ*metA* (Figure 2*E*). None of the supplemented metabolite concentrations resulted in complete strain loss, and all caused a net increase in combined co-culture growth rate (Figure 5*D-F*). We also found that the time between inoculation and steady state increased. This is likely due to the increase in the growth rate of both strains over the course of the entire experiment due to supplementation. While metabolite concentration changes are the main factor influencing growth rates in this system, their impact lessens at higher growth rates; the impact of the same relative increase in metabolite concentration on growth rate diminishes as concentrations and growth rates increase.

### Model captures experimental steady state

The cell population ratio in this system depends on factors such as the growth rates of ∼*metA* and Δ*argC* and rates at which they produce and consume methionine and arginine. To confirm our hypothesis about the mechanisms of ratio control and to predict the population ratio as a function of nutrient supplementation, we developed a mathematical model of the concentrations of Δ*metA* (*C*_*m*_), Δ*argC* (*C*_*a*_), arginine (*x*_*a*_) and methionine (*x*_*m*_) in the turbidostat, and fit the resulting model to population ratio data shown in Figure 2.

To construct our model, we assumed that Δ*metA* cells release excess arginine into the environment at a constant rate *β*_*a*_ and consume methionine at a rate that is proportional to the cell growth rate, with proportionality constant *η*_*m*_. We similarly assumed that Δ*argC* cells release methionine at rate *β*_*m*_ and consume arginine with proportionality constant *η*_*a*_. As shown in Figure 1*C* and *D*, the cell growth rates are increasing functions of metabolite. We therefore assumed that, after an initial adjustment period in the turbidostat, the dependence of cell growth rates can be modeled by Hill functions 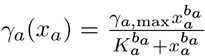 for Δ*argC* and 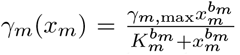 for Δ*metA*. Because the cells are grown in rich media prior to being added to the turbidostat, we hypothesize that the cells are in a different growth regime at the onset of the experiment. To account for this initial period, we assumed that the growth rate functions have the form *G*_*a*_(*x*_*a*_, *t*)= *ge*^−*at*^ + γ_*a*_(*x*_*a*_)(1 − *e*^−*at*^) for *argC* growth and *G*_*m*_(*x*_*m*_, *t*)= *ge*^−*at*^ + γ_*m*_(*x*_*m*_)(1 − *e*^−*at*^) for Δ*metA* growth in the model. The resulting model is nonautonomous and reflects the assumption that both cell types grow at a constant rate *g* at the beginning of the experiment, due to leftover nutrients from their preparatory growth. After adjusting to the turbidostat environment over approximately 1*/a* hours, the growth rates are modeled using Hill functions, *γ*_*a*_, and *γ*_*m*_.

To model dilution in the turbidostat, we also assumed a dilution rate function *δ* (*C*_*a*_, *C*_*m*_, *x*_*a*_, *x*_*m*_) defined so that the total cell concentration is constant, *C*_*a*_(*t*)+ *C*_*m*_(*t*)= *C*. Because methionine and arginine were supplemented in the dilution liquid, we included two source terms that represent supplementation concentrations *–* of arginine and *µ* of methionine added to the culture at a rate equal to the system dilution rate. We applied the conservation relation that is consistent with this dilution rate and scaled the concentration *C*_*m*_ by *C* to obtain an equation for the evolution of the population fraction 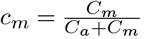. The resulting three-dimensional system of equations for *c*_*m*_ and the two metabolite concentrations has the form

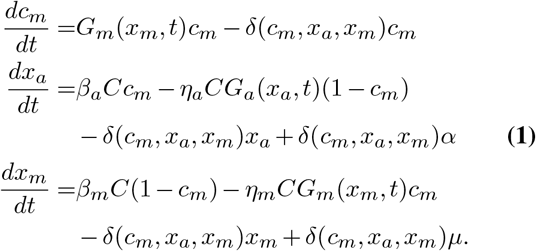

Further details about the model are provided in the Methods section, and a derivation of the nondimensionalized model can be found in the Supplemental Materials.

To ensure that the model captures the system response to supplemented nutrients, we fixed the supplementation parameters *α* and *µ* to the values used in the experiments shown in Figure 2, and obtained the remaining parameters by fitting the model to the 6 different population ratio datasets through a two-step Bayesian approach using the Metropolis-Hastings algorithm. The posterior parameter distributions are shown in Figure 6 and Figure 7. In Figure 3, we show the six datasets along with the population proportion, *c*_*m*_, obtained by solving the model equations numerically using parameter values sampled from the posterior distribution. The model correctly captures the steady state population ratios for all supplementation experiments. In Figure 8, we show the *c*_*m*_ trajectory obtained using posterior means, with the shaded region corresponding to one standard deviation of the inferred observational noise.

**Fig. 3.**
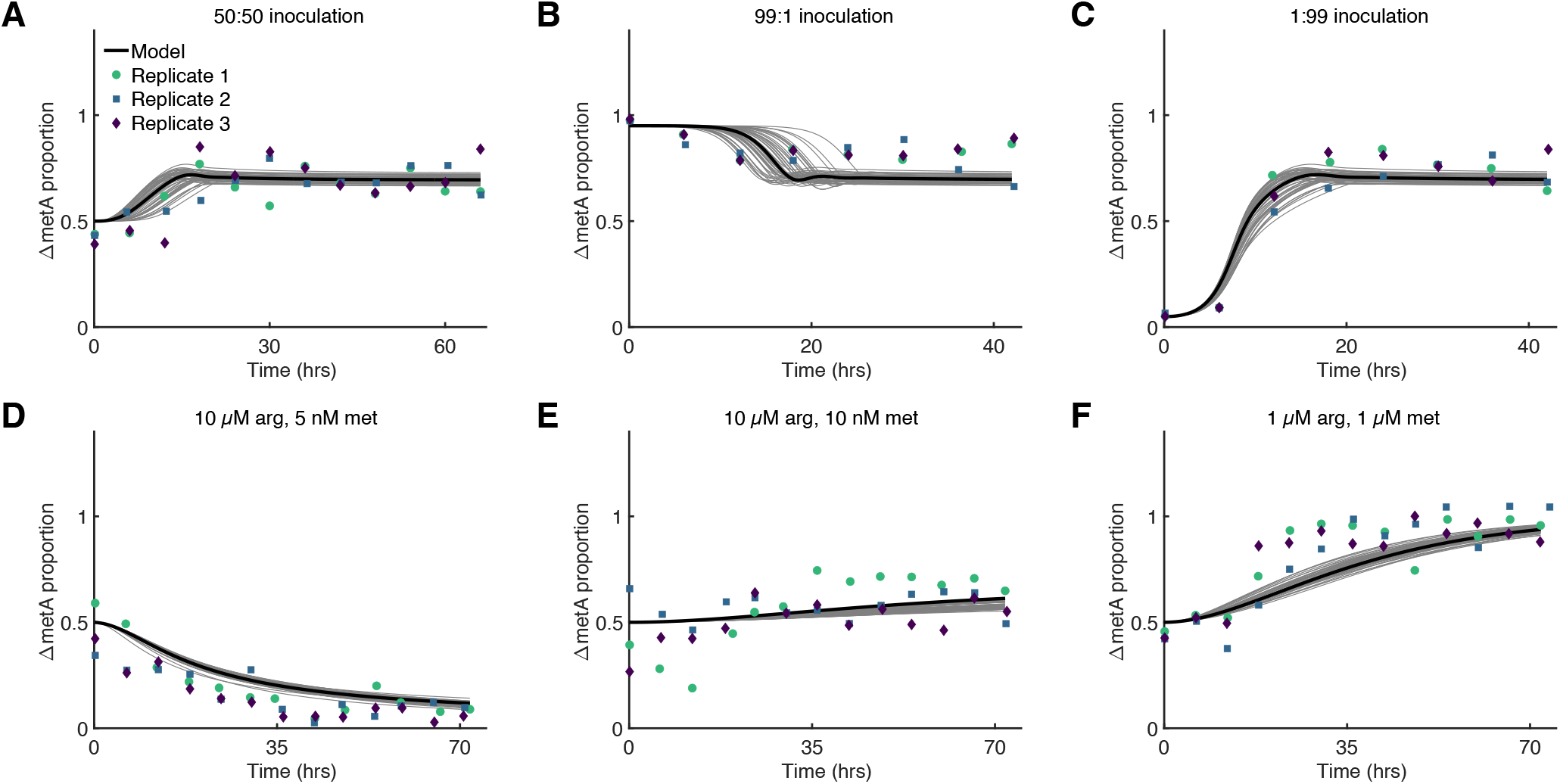
Mathematical modeling of turbidostat data. We used a Bayesian approach to fit our ODE model to the data shown in Figure 2. **(A-F)** Model solutions fit experimental data well. Solutions shown were generated using the mean parameter values (black curves) or samples (grey curves) from the posterior parameter distribution. Means of experimental proportions were calculated as in Figure 2 by summing the plate counts from each plate type and taking the summed ratio.

To confirm that the assumption of an initial transient in growth rates is needed, we fit an autonomous version of the model to data. To do so, we assumed that there is no transient period, and that growth rates are determined solely by the Hill functions, *γ*_*i*_(*x*_*i*_) for *i* ∈ {*a, m*}. Parameters and solutions for this autonomous model can be found in Figure 9 and Figure 10. When fit to data, this model produced solutions with strong damped oscillations that were absent in experimental data. Thus, the data suggest that including a transient growth state is necessary.

Parameter estimates suggest that the production rate of arginine is approximately an order of magnitude greater than the production rate of methionine. In both bulk culture growth rate experiments and supplementation experiments, more arginine is needed to increase the growth rate of Δ*argC* cells than methionine is needed to increase the growth rate of Δ*metA*. To increase the level of arginine in the model, we must either have a consistently high proportion of Δ*metA* cells, or a high arginine production rate. As the proportion of Δ*metA* cells is not always high, the model suggest that the arginine production rate must be larger than the methionine production rate. The growth rate Hill function parameters also indicate the relatively high concentration of arginine needed to increase Δ*argC* growth. The half maximum of the Δ*argC* cell growth rate must be about 130 times that of the Δ*metA* growth rate for the model to fit the data. This is a marked difference from the cell growth rates in bulk culture, where the half maxima of the growth rate curves are within an order of magnitude of one another. The inferred maximum cell growth rates are similar (*γ*_*m*,max_ = 1.014 *γ*_*a*,max_), implying similar growth rates for the strains when nutrients are present in abundance. Moreover, the posterior distribution for the parameter ratio 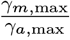 has a very small variance, confirming our hypothesis that the cell ratio *c*_*m*_ depends strongly on the balance of growth rates of the two cell types.

### Predictions and model validation

We next asked how well our model can predict experimentally observed strain ratios. To do so we first obtained steady state ratio prediction over a range of supplementation concentrations, *µ* and α. Figure 4*A* shows that the model predicts the population ratio can be adjusted by varying arginine supplementation between 0 *µ*M and approximately 1.5 *µ*M, and methionine supplementation between 0 nM and approximately 15 nM.

**Fig. 4.**
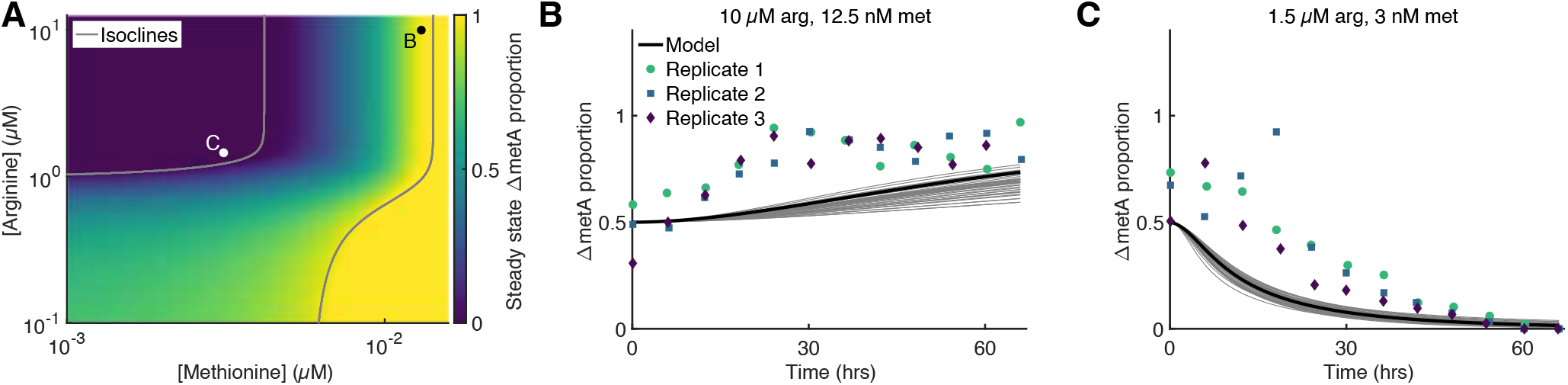
Model prediction and validation. **(A)** Heatmap of predicted steady state proportions of Δ*metA* at different arginine and methionine concentrations. Ratios were obtained by finding steady state solutions of Eq. Eq. (1). Dots indicate predictions selected for experimental validation, with labels indicating the relevant panel. Lines enclose regions in parameter space where one strain is predicted to be extinct. **(B)** and **(C)** Predictions of the model compared to experimental data. Numerical solutions were generated using the mean parameter values (black curves) or random samples (grey curves) from the posterior parameter distributions. Means of experimental proportions were calculated as in Figure 2 by summing the plate counts from each plate type and taking the summed ratio. Both supplementation values moderately increased the overall co-culture growth rate (Figure 11).

**Fig. 5.**
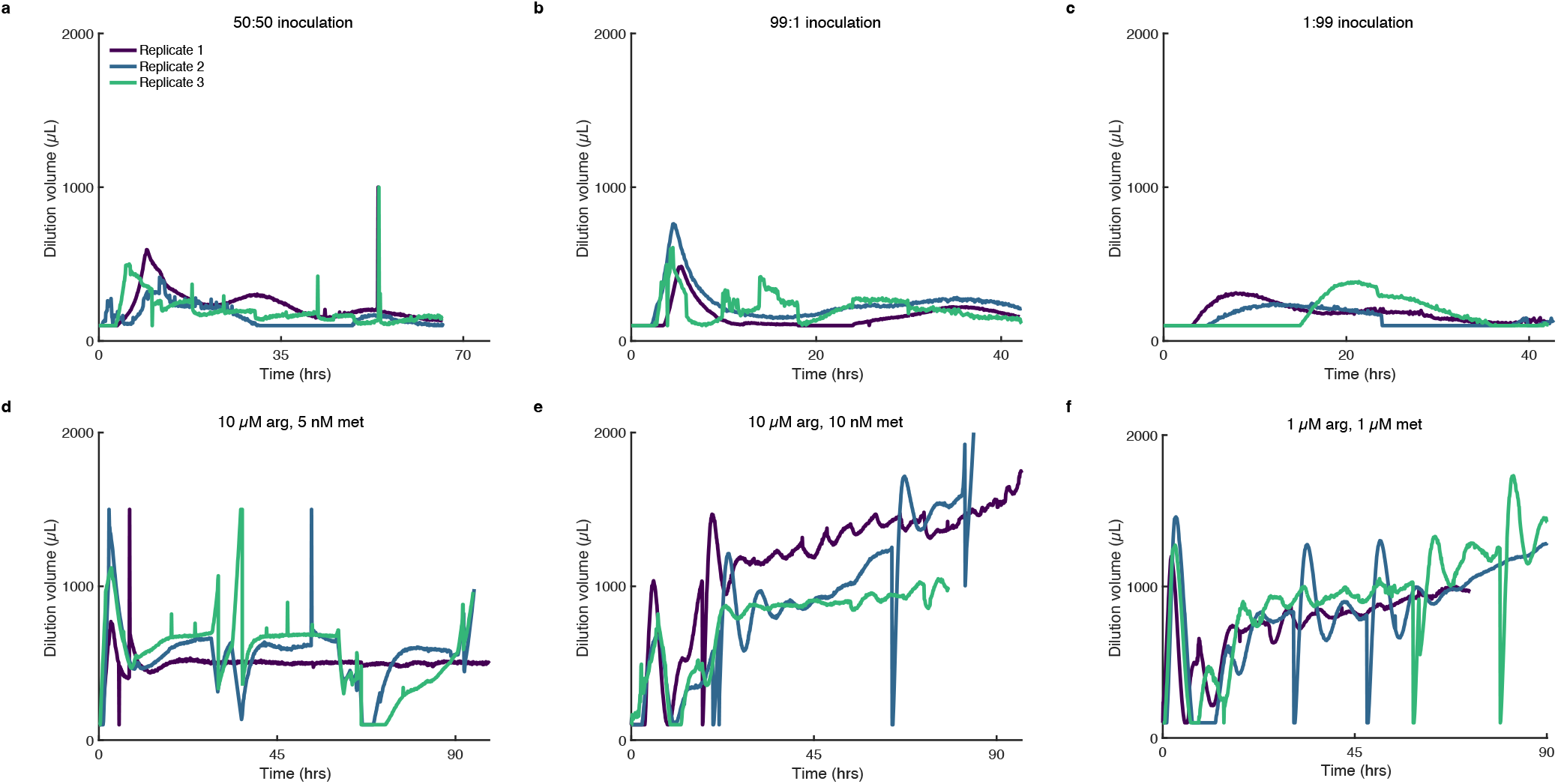
Turbidostat dilution volumes during coculture experiments. At each time step, the turbidostat estimates the current OD, compares it to the setpoint OD (0.2 in all experiments), and uses a proportional integral controller to determine the appropriate dilution volume to add to return the coculture to the setpoint OD. Dilution plots correspond to ratio plots in Figure 2. Spikes are due to misreads from OD chamber.

We tested these predictions in two supplementation experiments. For the first experiment (Figure 4*B*), we chose 10 *µ*M arginine and 12.5 nM methionine, where the model predicts a steady state ratio of approximately 0.9. Achieving this ratio would experimentally confirm that the system can maintain a high proportion of Δ*metA* without eliminating Δ*argC*. For the second experiment (Figure 4*C*), we chose 1.5 *µ*M arginine and 3 nM methionine, because the model predicts that Δ*argC* cells will take over the culture - a result we did not see in our initial experiments (Figure 2). In both cases, experimentally observed ratios at steady state agreed well with model predictions.

## Discussion

In this work, we demonstrated that mutualistic cross-feeding in *E. coli* is a powerful tool for controlling the population abundances in a two-strain co-culture. While we used a pair of methionine and arginine auxotrophs, we believe that this tight control mechanism can be extended to other auxotrophic combinations that have been shown to grow mutualistically in minimal media (33, 34). Indeed, previous results have demonstrated multi-population maintenance in larger synthetic consortia containing mutualistic auxotrophs (35, 40), suggesting that such tight control may extend to consortia with more than two strains.

Mutually auxotrophic ratio control depends on a lack of competition for other nutrient resources. In regimes with limited carbon sources, for example, the concentration of arginine and methionine would no longer have a dominant effect on the growth rate of each strain. While both strains could remain in regimes without any supplemented amino acids, tuning via supplementation would be much more difficult, as the relative range of non-competitive growth rates for each strain would be narrower.

Our mathematical model lends further support for the proposed mechanism of ratio control in this system. The model relies on simplifying assumptions about cell metabolism, mutualism, and dilution to a finite population size: We assumed that cells are initially in a constant growth rate regime determined by the conditions they experienced before transfer to the turbidostat. Upon transfer, cells shift to a regime where their growth rates depend on the nutrient they need to survive. The data support this assumption. However, a more detailed understanding of the metabolic mechanisms that determine growth in these two regimes requires further study. Additionally, our model relies on the assumption that the turbidostat continuously dilutes the culture to keep the cell concentration constant. In reality, dilutions occur every 5 minutes. These increments are likely small enough compared to the timescale of changes in ratio, so that our use of continuous dilution in our model is justified. Modeling populations that experience dilutions less frequently would require a different dilution term. Even with these simplifying assumptions, fitting our model to data gave us insight into how cells grow and exchange nutrients in the turbidostat. Additionally, the model predicted the outcome of supplementing nutrients at a range of concentrations.

This work presents new tools for controlling and understanding population abundances in microbial consortia. While work remains to be done to demonstrate its compatibility with other engineering practices, such as classical metabolic engineering or other forms of consortia control, the ability to supplement these cultures with growth-enhancing metabolites without breaking ratio control reduces the toxicity risks seen when using other mechanisms. Overall, its simplicity and predictability make it a promising candidate for future advances in synthetic microbial consortia.

## Materials and Methods

### *E. coli* strains

All experiments were conducted in the Δ*argC* and Δ*metA* strains taken from the Keio collection (41) (parent strain *E. coli* K-12 BW1125), which both contain a kanamycin resistance gene in place of their deletion.

### Growth media

In all experiments, both Δ*argC* and Δ*metA* monocultures were first grown up overnight in LB medium containing kanamycin (kan; 50 mg/L). co-culture experiments were conducted in minimal M9 media, comprised of 1X M9 salts, 0.1 mM CaCl_2_, 1 mM MgCl_2_, 0.4% glucose, and 50 mg/L kanamycin (43). Arginine and methionine were added in relevant assays at indicated concentrations during media prep.

### Growth rate assay

Overnight auxotroph cultures were grown for 1 hour in rich media and serially diluted in M9 media to equivalent ODs, calculated at 0.005. These were then loaded into 96-well plates and grown in a Tecan Spark plate reader with incubation (37C) and shaking (216 rpm) with measurements taken every 10 minutes for 24 hours. Growth rates were then estimated by fitting the background subtracted OD data to a logistic growth curve in MATLAB using the `lsqcurvefit` function. See Supporting Information for more details.

### Turbidostat

The turbidostat used in this work was adapted from a version created by the Klavins lab (42, 44). Briefly, our turbidostat follows the design which can be followed at https://depts.washington.edu/soslab/turbidostat/pmwiki/, with the main exception being that we used a Raspberry Pi to control the behavior of the turbidostat, rather than a full computer, which further reduced the cost and enabled the building of multiple turbidostats in the same lab space.

### Turbidostat assay

Overnight auxotroph cultures were grown for 1 hour in rich media and then mixed at equi-OD ratios, unless otherwise indicated. These were then loaded into the culture tube via syringe, which allowed for penetrating the rubber stopper without contaminating the culture tube. Samples were taken by collecting effluent into a micro-centrifuge tube every 6 hours. Samples were serially diluted into M9 media in 1:10 dilutions 4 times, with at least 10 pipette mixing steps in between each dilution to minimize dilution variance. Samples were then added to both a rich media kanamycin plate and a minimal M9 media kanamycin plate supplemented with 100 *µ*M methionine. This plating was done by adding 10 *µ*L of diluted sample as a spot. The plate was then tilted to allow the spot to streak down the plate due to gravity, to avoid cell loss that would potentially result from a tactile spreading mechanism. Plates were grown at 37C either overnight (rich media) or for 48 hours (minimal media). Colonies were then counted and recorded by hand. Plate counts and turbidostat recordings can be found in datasets S1 and S2.

### Mathematical model

The full model consists of a system of four ordinary differential equations that describe how the concentrations (in *µ*M) of the following species change over time (in hours): Δ*argC* (*C*_*a*_), Δ*metA* (*C*_*m*_), arginine (*x*_*a*_) and methionine (*x*_*m*_). The resulting ordinary differential equations have the form:

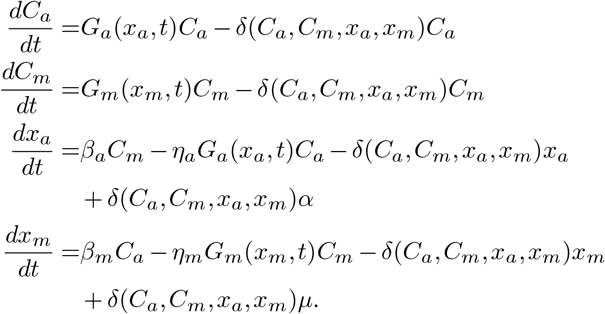

The dilution term, *δ*(*C*_*a*_, *C*_*m*_, *x*_*a*_, *x*_*m*_), is defined so that the total cell concentration remains constant, *C*_*a*_ + *C*_*m*_ = *C*, or 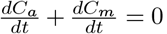. Plugging in the definitions of 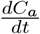 and 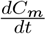 and setting their sum to 0 gives,

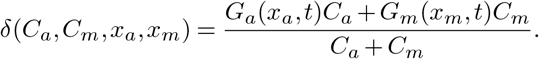

As there are billions of cells in the turbidostat, we assume that stochastic fluctuations can be neglected. The turbidostat dilutes the culture every 5 minutes, but we assume that these intervals are sufficiently small compared to the timescale of changes of the different concentrations so that dilution can be modeled as occurring continuously in time. The conservation relation, *C*_*a*_ + *C*_*m*_ = *C*, also allows us to reduce this system to 3 equations, and scale the *C*_*m*_ equation by *C* to obtain Eq. (1).

### Parameter fitting

We used the DEMetropolisZ sampler in the PyMC Python library (46) to fit our model to data. We assumed that the cells take about 15 hours to shift to the Hill Function growth regime in the turbidostat (based on observations from the turbidostat dilution rates), so we fit the model parameters in two steps: First, we fit the autonomous model to data collected at time points greater than 15 hours in datasets shown in Figures 2 A, D, E, and F. Second, we fit the two remaining parameters of the nonautonomous model to data collected at time points between 0 and 15 hours in the six datasets in Figure 2, while setting the remaining parameters equal to the mean of the posterior in the first step. Code and data are accessible at github.com/amandaalexander/AuxotrophicCrossFeeding.

## ACKNOWLEDGEMENTS

This work was supported by funding from the joint National Science Foundation and National Institutes of Health Mathematical Biology Program grant 1R01GM144959 (K.J., M.R.B.), and the National Science Foundation grants MCB-1936774 (M.R.B), and MCB-1936770 (K.J.). A.M.A supported by a training fellowship from the Gulf Coast Consortia, on the NLM Training Program in Biomedical Informatics & Data Science (T15LM007093).

## Supplementary Note 1: Model description

To model the dynamics of the consortium in a turbidostat, we developed a model consisting of a system of ordinary differential equations. The model describes the change in concentrations (in micromolar) of the following species over time (in hours): Δ*argC* (*C*_*a*_), Δ*metA* (*C*_*m*_), arginine (*x*_*a*_) and methionine (*x*_*m*_). Because there are billions of cells in the turbidostat, we ignore stochastic fluctuations in cell population levels and assume that the concentration of all cell and molecular species can be modeled as continuous functions of time. Additionally, we assume that the contents of the turbidostat are well mixed.

The model incorporates cellular production and consumption of metabolites. Δ*argC* cells secrete excess methionine at constant rate *β*_*m*_ and metabolize arginine from their environment at a rate proportional to their growth rate, with proportionality constant *η*_*a*_. Similarly, Δd*metA* cells secrete excess arginine at constant rate *β*_*a*_ and metabolize methionine at a rate proportional to their growth rate with proportionality constant *η*_*m*_. Biologically, Δ*argC* and Δ*metA* cells use some of the methionine and arginine that they respectively produce, but we exclude this from the model and hence interpret *β*_*m*_ and *β*_*a*_ as rates of metabolite production in excess of what is consumed by the producing cells.

All molecule and cell species are diluted out of the system by the turbidostat at a rate *δ(C*_*a*_, *C*_*m*_, *x*_*a*_, *x*_*m*_) that maintains a constant turbidity, which we assume implies a constant total concentration of cells. We define the dilution rate, in units of hours^−1^, to enforce the conservation relation *C*_*a*_ + *C*_*m*_ = *C*, where *C* is constant. Experimentally, exogenous arginine and methionine are introduced into the system via the dilution liquid. This is modeled by source terms *αδ*(*C*_*a*_, *C*_*m*_, *x*_*a*_, *x*_*m*_) and *µδ*(*C*_*a*_, *C*_*m*_, *x*_*a*_, *x*_*m*_) in the arginine and methionine equations, respectively. Here *α* is the concentration of arginine and *µ* is the concentration of methionine.

Each auxotrophic cell population in the model grows at a rate that depends on the concentration of the metabolite that the population cannot produce. We consider two possible forms for these growth rates. First, we consider growth rate functions *γ*_*i*_(*x*_*i*_), for *i* ∈ {*a, m*}, that have the form

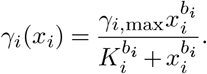

The choice of Hill functions reflects the experimental observation that cells grow at a saturating growth rate when provided with excess nutrients, but that growth rate is lower as nutrient concentrations decrease. This observation was made with cells in bulk culture, and we assume that the general principle that nutrients increase cell growth rates holds in the turbidostat. However, we do not assume that the parameter values of these Hill functions, *i*.*e*. dependence on nutrient concentration, is the same as in bulk culture.

Using growth rate functions, *γ*_*i*_(*x*_*i*_), that are not explicitly dependent on time is equivalent to assuming that cells are in the same metabolic regime during the entire experiment. However, right after the initial transfer to the turbidostat cells are likely to carry stored nutrients, and thus the initial dependence of their growth rate on the ambient concentration of nutrients is likely to be different: Stored nutrients will decrease the growth rate dependence on nutrient concentration in the turbidostat during the first several hours of the experiment, with a gradual transition to the growth rate dependence modeled by Hill Functions, *γ*_*i*_(*x*_*i*_). Thus, the second form of growth rate functions we consider have the form

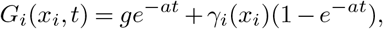

where *g* is a constant representing the initial growth rates and *a* represents the inverse of the timescale at which the cells shift between the two growth regimes. With these growth rates, our model becomes nonautonomous, but solutions converge to the same steady state value as the autonomous version.

The resulting model has the form:

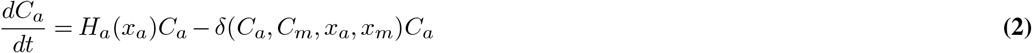

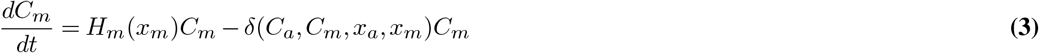

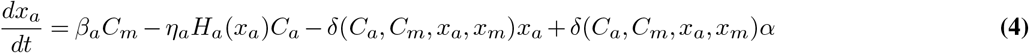

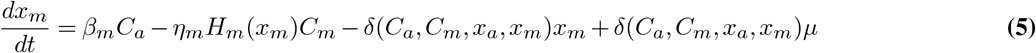

where *H*_*i*_(*x*_*i*_), for *i* ∈ {*a, m*}, equals either *γ*_*i*_(*x*_*i*_) or *G*_*i*_(*x*_*i*_, *t*). The dilution term *δ*(*C*_*a*_, *C*_*m*_, *x*_*a*_, *x*_*m*_) has the form

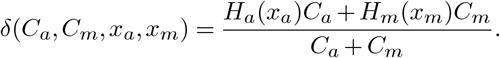

This form for *δ*(*C*_*a*_, *C*_*m*_, *x*_*a*_, *x*_*m*_) enforces the assumption that *C*_*a*_ + *C*_*m*_ = *C*, or 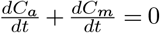

## Supplementary Note 2: Steady state analysis

For steady state analysis, it is useful to consider an ODE for the fraction 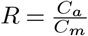. We use the quotient rule to find:

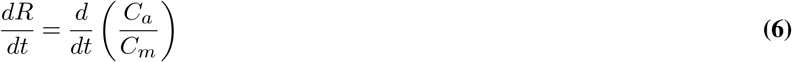

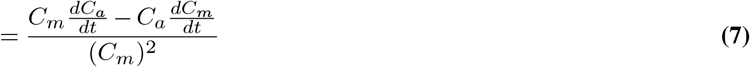

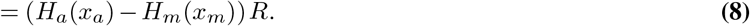

The quantities on each side of this equation have units of *t*^−1^. We are interested in the case where *R>* 0 and *R<* ∞, because this represents coexistence of the two cell strains. If we assume the ratio *R* is finite and nonzero, then to achieve steady state we must have 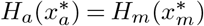, at steady-state metabolite concentrations 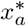 and 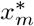. Thus the model makes the reasonable prediction that for the cell strains to coexist at steady state, the steady state metabolite concentrations must be such that the growth rates of the two strains are equal.

## Supplementary Note 3: Nondimensionalization

We nondimensionalized the model for simplicity. Here we also discuss which parameters are known and which parameters we fit to data from turbidostat experiments. We focus on the model with autonomous growth rate functions, *γ*_*i*_(*x*_*i*_), as the nondimensionalization is similar for the non-autonomous case.

### Autonomous ODEs

The dimensional equations are

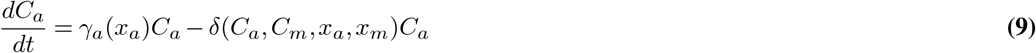

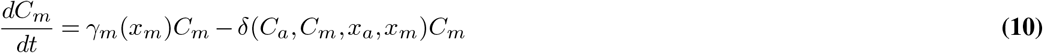

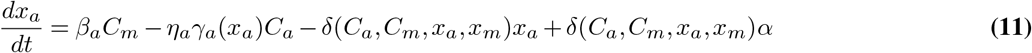

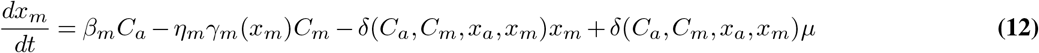

where

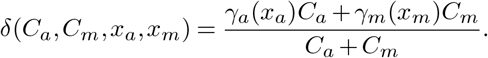

We define nondimensional time parameter *τ* as *τ* = *γ*_*a*,max_*t*, where *γ*_*a*,max_ is the maximum value of the Δ*argC* growth rate function *γ*_*a*_(*x*_*a*_). To nondimensionalize the metabolite equations, we divide by *K*_*m*_, the half max constant of the Δ*metA* growth rate function *γ*_*m*_(*x*_*m*_). To nondimensionalize the cell concentration equations, we take advantage of the conservation relation *C*_*a*_ + *C*_*m*_ = *C* and measure cell concentrations relative to the constant concentration, *C*. We have three resulting equations for 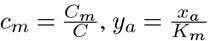 and 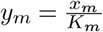:

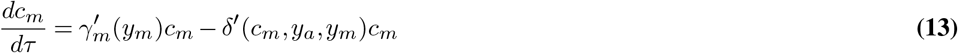

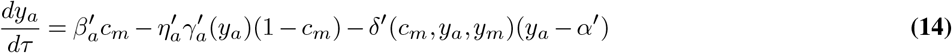

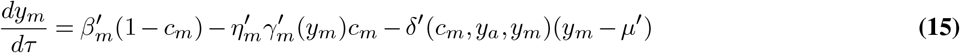

where the growth rates are 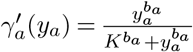 and 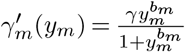, and the dilution function is 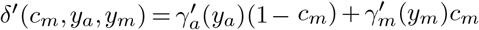.

We have eight resulting nondimensional parameters that we will fit to data:

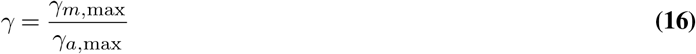

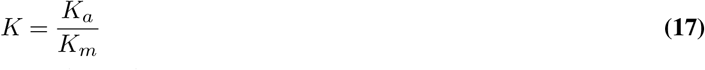

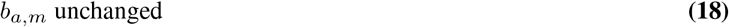

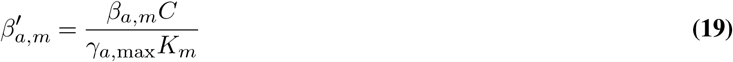

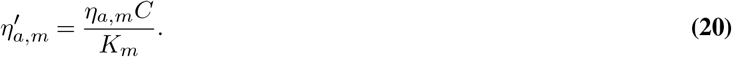

Additionally, the nondimensional supplementation parameters are 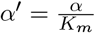 and 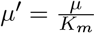. Since we controlled the level of supplementation, when fitting the model to data, we assumed that these parameters were determined by the supplementation concentration in each experiment.

### Nonautonomous ODEs

In the case of the nonautonomous ODEs with growth rates

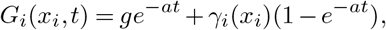

we nondimensionalize by rescaling the equations in the same way as above. The resulting nondimensional growth rates are 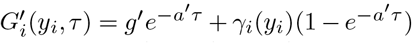. The nonautonomous model has the same 8 nondimensional parameters as the autonomous model, and two other parameters: 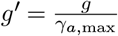 and 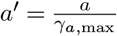.

## Supplementary Note 4: Parameter inference

Because we wish to quantify the uncertainty in the model parameters, we use Bayesian inference to fit the model to six cell ratio timecourse datasets.

Before all fitting procedures, we process the data from the six turbidostat experiments in the following way:

- Take the average of the three technical replicates of plating on LB during each experiment, and the three technical replicates of plating on minimal media during each experiment
- Use these average values to calculate ratio data for each of the three biological experiments
- If a data point indicates a ratio greater than 1, set that point equal to 1
- nondimensionalize the times at which samples were taken in the same way we nondimensionalized the model: by multiplying times by *γ*_*a*,max_

The model dynamics depend on the parameter ratios, 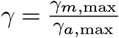 and 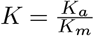, and not individual parameters. However, we assume that *γ*_*a*,max_ = 2.3*/*hr for the purpose of nondimensionalizing the data. Biologically, we are assuming that the cells can divide at a maximum rate of a little more than twice per hour given all the nutrients they need. This is in line with our knowledge of *e*.*coli* growth rates.

Similarly, we need to nondimensionalize the values representing supplemented methionine and arginine concentrations, for our nondimensionalized model to replicate our experimental modulation of methionine and arginine. Because we divided the metabolite equations in our model by *K*_*m*_ in our nondimensionalization, we divide our experimental supplementation concentrations by *K*_*m*_ as well. To obtain numerical values, we assume that *K*_*m*_ = 0.006*µ*M. This is an estimate based on Figure 2d and e: when we fix our arginine supplementation concentration, changing our methionine supplementation concentration from 5nM to 10nM augments the steady state ratio significantly. We therefore conclude that a value of approximately 6nM methionine allows cells to grow at half their maximum growth rate.

In the following sections, we describe our procedure for fitting the parameters of the autonomous model, and then our procedure for fitting the parameters of the nonautonomous model. The autonomous model cannot capture the dynamic phase of the cell ratio timecourse data as well as the nonautonomous model. Thus, the data suggest that the cells start in a different metabolic regime than they are in at steady state.

### Autonomous model

We fit the autonomous model simultaneously to the six experimental datasets for the cell ratio *c*_*m*_ measured as a function of time, and shown in Figure 2. To do this, we use the DEMetropolisZ sampler in PyMC, a Python package for Bayesian statistical modeling. We use 10000 tuning steps to avoid mixing issues. We draw 10000 samples from 4 chains after the tuning period.

The initial conditions of the model were not fit to data: We set the initial ratio *c*_*m*_(0) to 0.5 to reflect the supplemented experiments, and 0.95 and 0.05 to reflect the respective experiments with 0 supplementation and varying inoculation ratio. We use initial conditions *y*_*a*_(0) = 150 and *y*_*m*_(0) = 150 because cells were grown in rich media before the start of the turbidostat experiment and therefore have nutrient stores at time 0 of the experiment. Changing the numerical value of 150 does not have a large impact on the results. The purpose of using these high initial conditions is to give the autonomous model the best chance at capturing the effect of the initial nutrient stores in the cells without explicitly modeling a different growth regime at the start of the experiments.

We use the priors found in Table 1. We selected the priors for *β*_*a*_, *β*_*m*_ based on our knowledge that the system is much more sensitive to low concentrations of methionine supplementation than arginine supplementation. We assumed that *K* is relatively high following the same reasoning. We selected low priors for *η*_*a*_ and *η*_*m*_ because the metabolite consumption rates must not be greater than the production rates. Our prior for *γ* reflects the assumption that it is unlikely that one cell species grows twice as fast as the other. Finally, we chose uninformative priors for the Hill coefficients *b*_*a*_ and *b*_*m*_.

**Table 1.** Priors used when fitting autonomous model to data.

| Parameter | Prior |
| --- | --- |
| $\beta_a$ | Gamma(mean=100, sigma=50) |
| $\beta_m$ | Gamma(mean=1, sigma=1) |
| $\eta_a$ | Gamma(mean=0.1, sigma=0.2) |
| $\eta_m$ | Gamma(mean=0.1, sigma=0.2) |
| $\gamma$ | Uniform(lower=0.5, upper=2) |
| $K$ | Gamma(mean=700, sigma=250) |
| $b_a$ | Uniform(lower=0.5, upper=10) |
| $b_m$ | Uniform(lower=0.5, upper=10) |

The posterior distributions are shown in Figure 9. In Figure 10, we used the posterior mean to generate the predicted timecourse of the ratio, and plot it along with the experimental data. The model captures the steady state of the system well for all supplementation concentrations, but the model does not capture the transient phase well. Thus, we fit the nonautonomous model to the data to see if the data is better explained by a model that includes a separate metabolic regime in the beginning of the experiments.

### Nonautonomous model

To ensure the parameters are constrained by the data, we fit the model in two phases. In both phases, we used DEMetropolisZ sampler in PyMC. We used the same initial conditions for *c*_*m*_ as in the autonomous model fit, but for *y*_*a*_ and *y*_*m*_ we used initial conditions equal to the respective metabolite supplementation concentration in each experiment. We did so because the nonautononomous growth rates *G*_*i*_(*x*_*i*_, *t*) account for the initial stored metabolites.

#### Phase 1

We fit the autonomous model to data shown in Figure 2, only using measurements at times greater than 15 hours. Because the nonautonomous model converges to the autonomous model over time, this method allows us to constrain all parameters except for *a* and *g* in the nonautonomous model. This approach also ensures that the model parameters are such that the model captures the observed steady state cell ratios. In this fit, we use 6000 tuning steps and draw 6000 samples from 4 chains after the tuning period. We used the priors given in Table 2. The posterior distributions are shown in Figure 6. We choose the mean for the parameter values used in the the main body of the text.

**Table 2.** Priors used when fitting nonautonomous model to data.

| Parameter | Prior | Phase |
| --- | --- | --- |
| $\beta_a$ | Gamma(mean=100, sigma=50) | 1 |
| $\beta_m$ | Gamma(mean=1, sigma=1) | 1 |
| $\eta_a$ | Gamma(mean=.1, sigma=.2) | 1 |
| $\eta_m$ | Gamma(mean=.1, sigma=.2) | 1 |
| $\gamma$ | Uniform(lower=0.5, upper=2) | 1 |
| $K$ | Gamma(mean=700, sigma=250) | 1 |
| $b_a$ | Uniform(lower=0.5, upper=10) | 1 |
| $b_m$ | Uniform(lower=0.5, upper=10) | 1 |
| $a$ | Uniform(lower=.01, upper=.15) | 2 |
| $g$ | Uniform(lower=1, upper=2) | 2 |

**Fig. 6.**
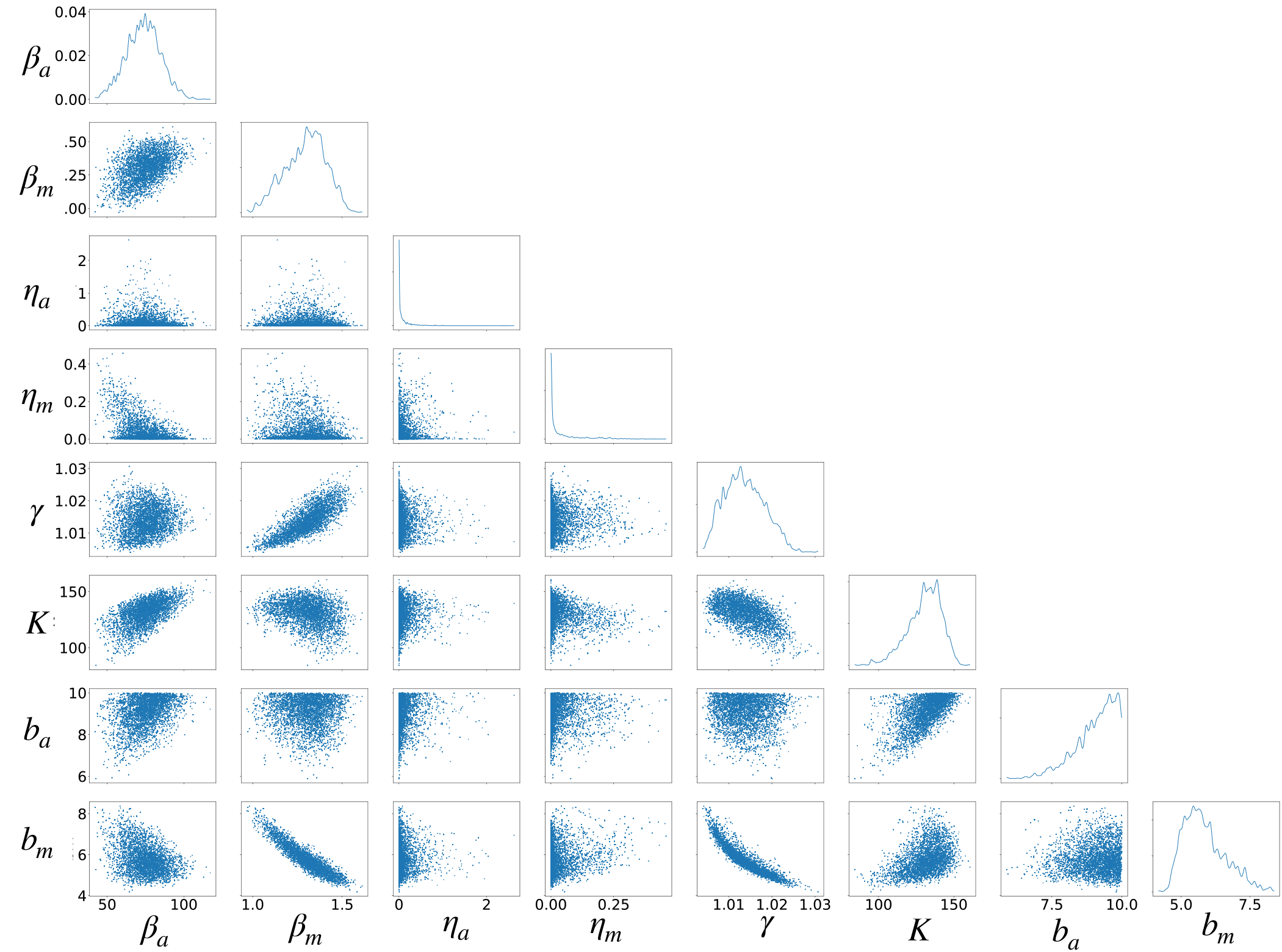
Result from using DEMetropolisZ sampling to fit the autonomous model to experimental data after an initial period of 15 hours. We show marginals from the posterior distribution of parameter values. We use the mean of this posterior to fix the parameter values of the nonautonomous model and subsequently fit *a* and *g* to the experimental data between 0 and 15 hours of the experiments.

**Fig. 7.**
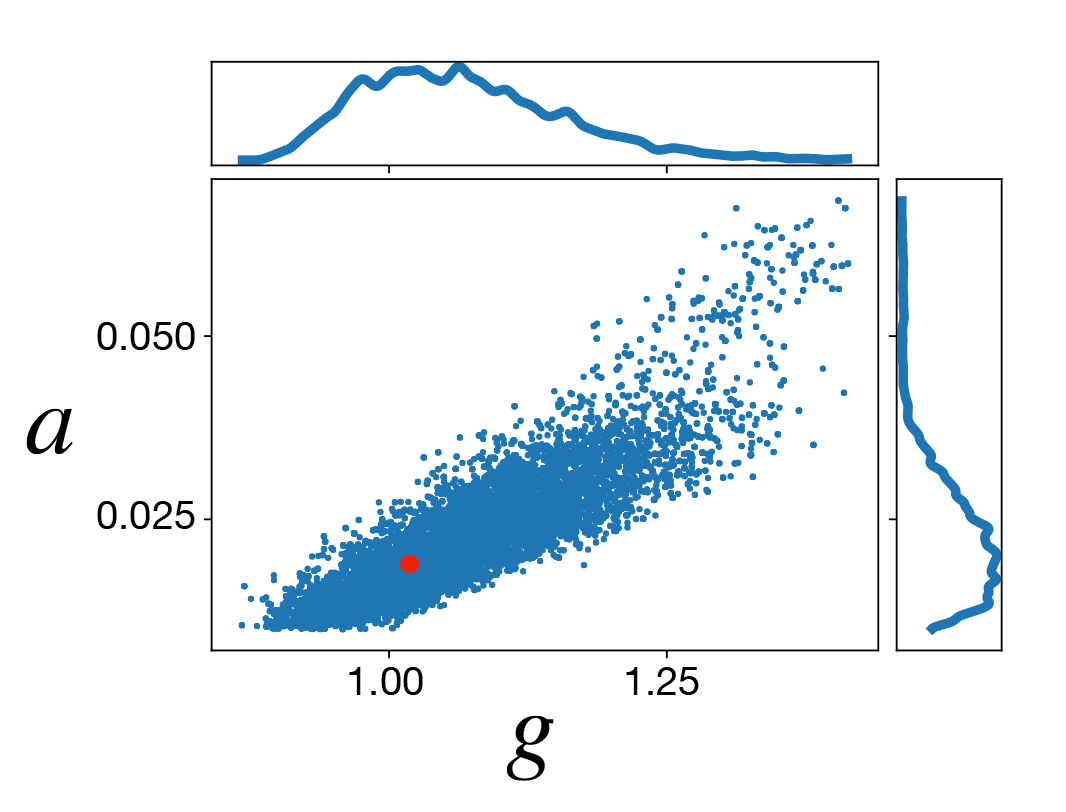
Result from using DEMetropolisZ sampling to fit the nonautonomous model parameters *a* and *g* to experimental datapoints between hours 0 and 15. We show marginals from posterior distribution of parameter values. There is a correlation between *a* and *g*, and we select (*g, a*)= (1.068, 0.022) (marked with red point) for the results in the main body of the text.

**Fig. 8.**
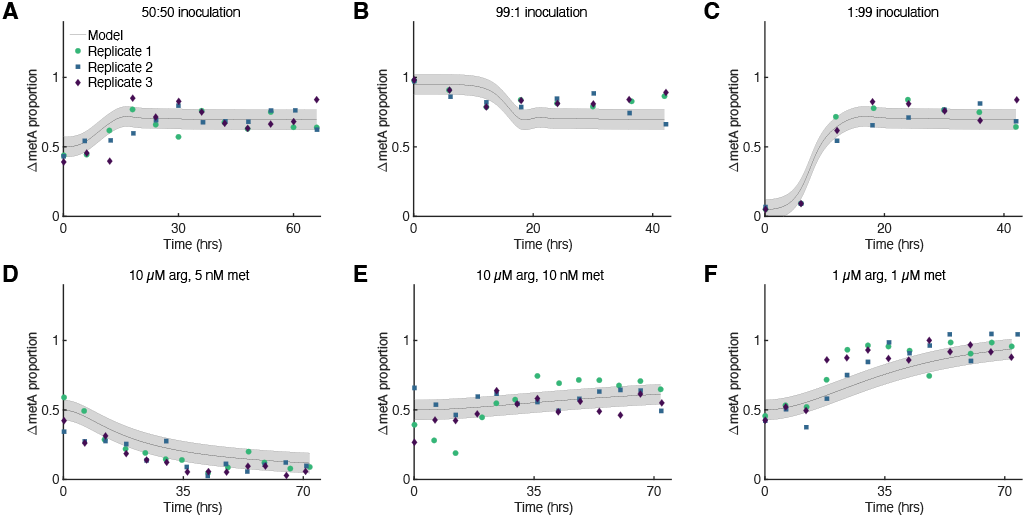
On the same axes as experimental ratio data, we plot the mean parameter set from the posterior in Figure 6 with (*g, a*)= (1.068, 0.022). The grey shaded region depicts the mean ratio plus and minus one standard deviation in the observational noise, as predicted by DEMetropolisZ. The horizontal axis is nondimensionalized time *τ*.

**Fig. 9.**
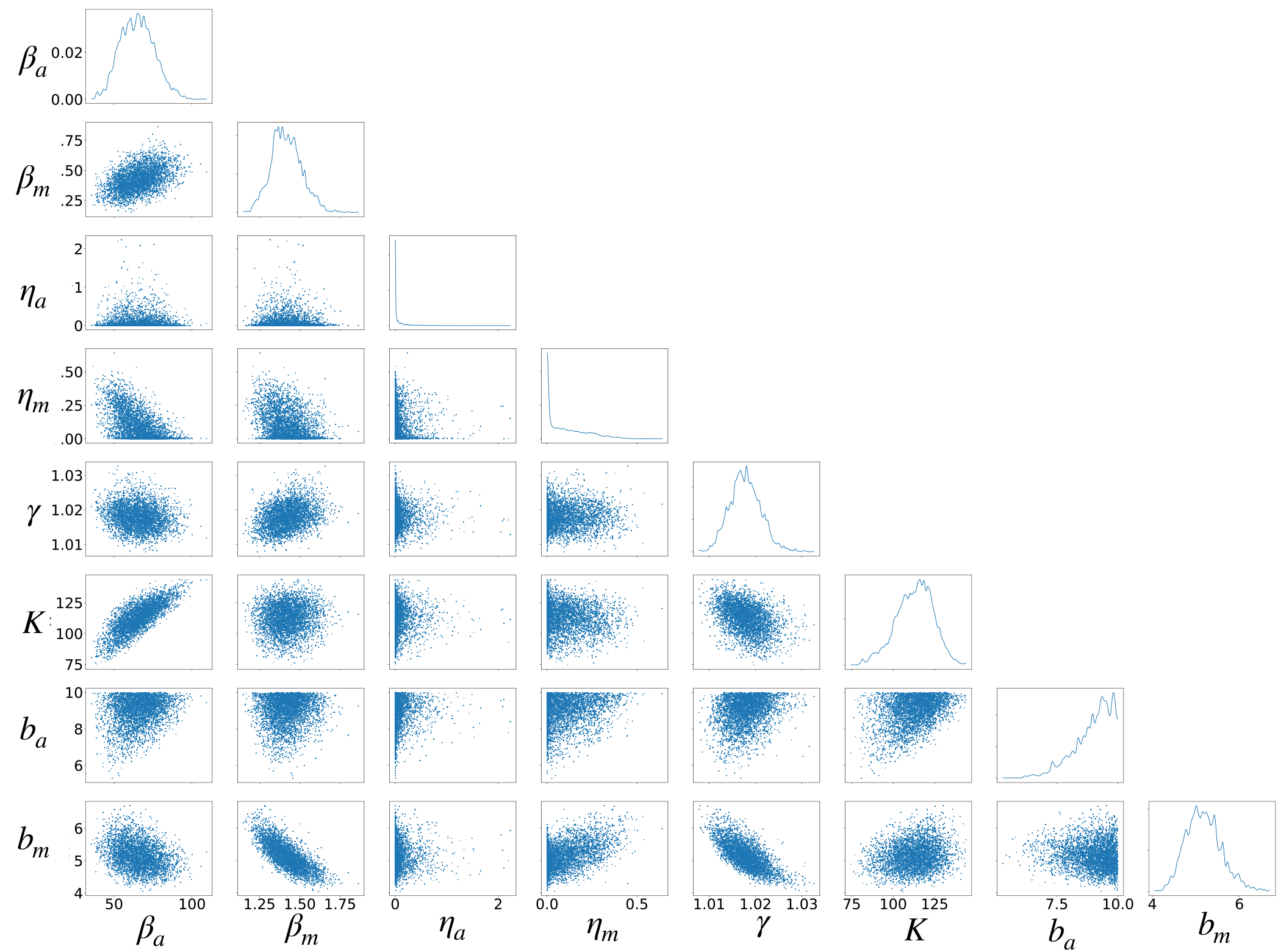
Result from using DEMetropolisZ sampling to fit the autonomous model to experimental data. We show marginals from posterior distribution of parameter values.

**Fig. 10.**
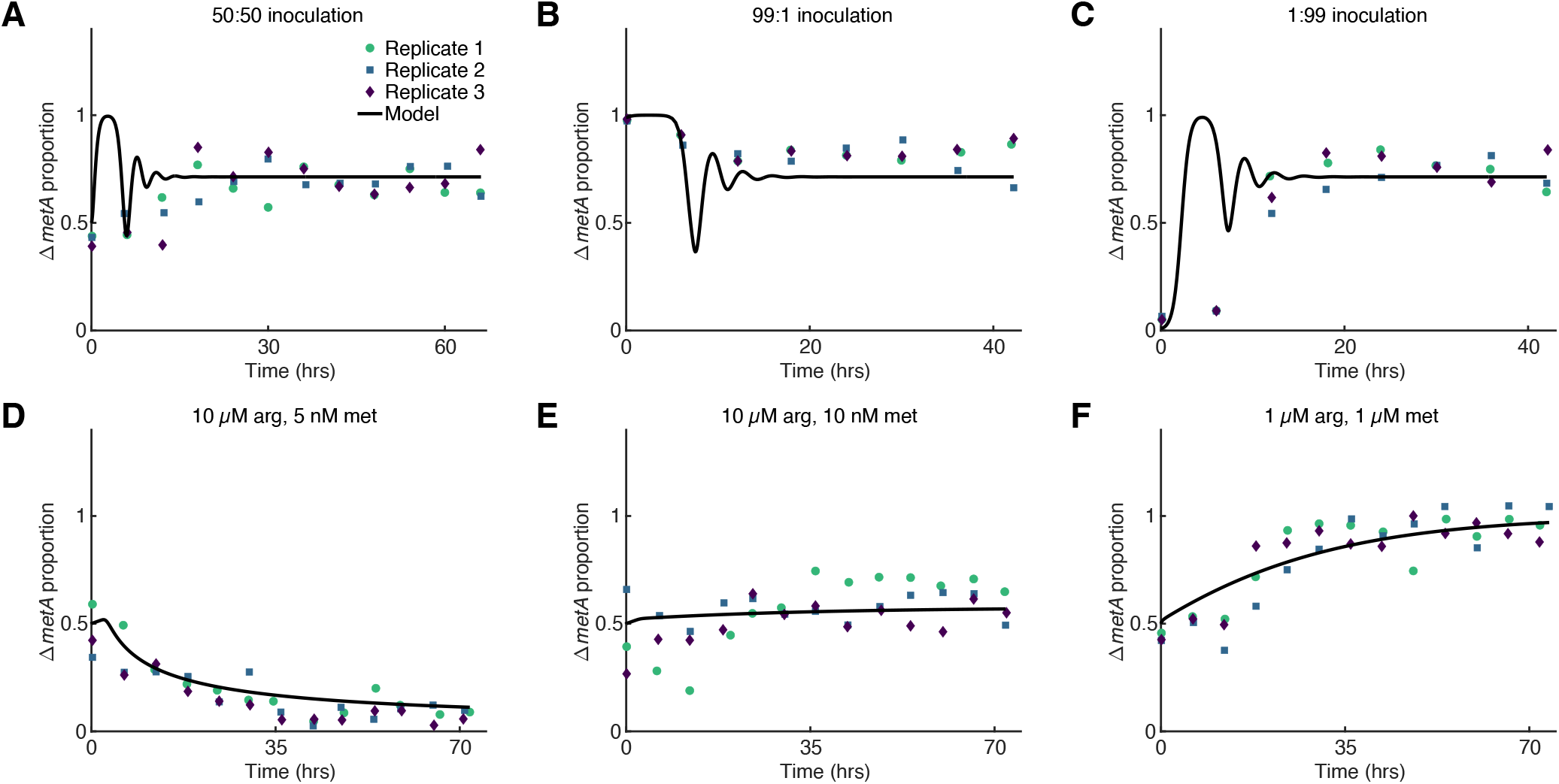
Results of autonomous mathematical model of turbidostat data. We fit our autonomous ODE model to our experimental data shown in Figure 2. **(A-F)** Model solutions fit experimental steady state data well, but the model solutions have initial damped oscillations that are not seen in the data. Solutions were generated using the mean of the posteriors obtained via DEMetropolisZ sampling. Means of experimental proportions were calculated as in Figure 2 by summing the plate counts from each plate type and taking the summed ratio.

**Fig. 11.**
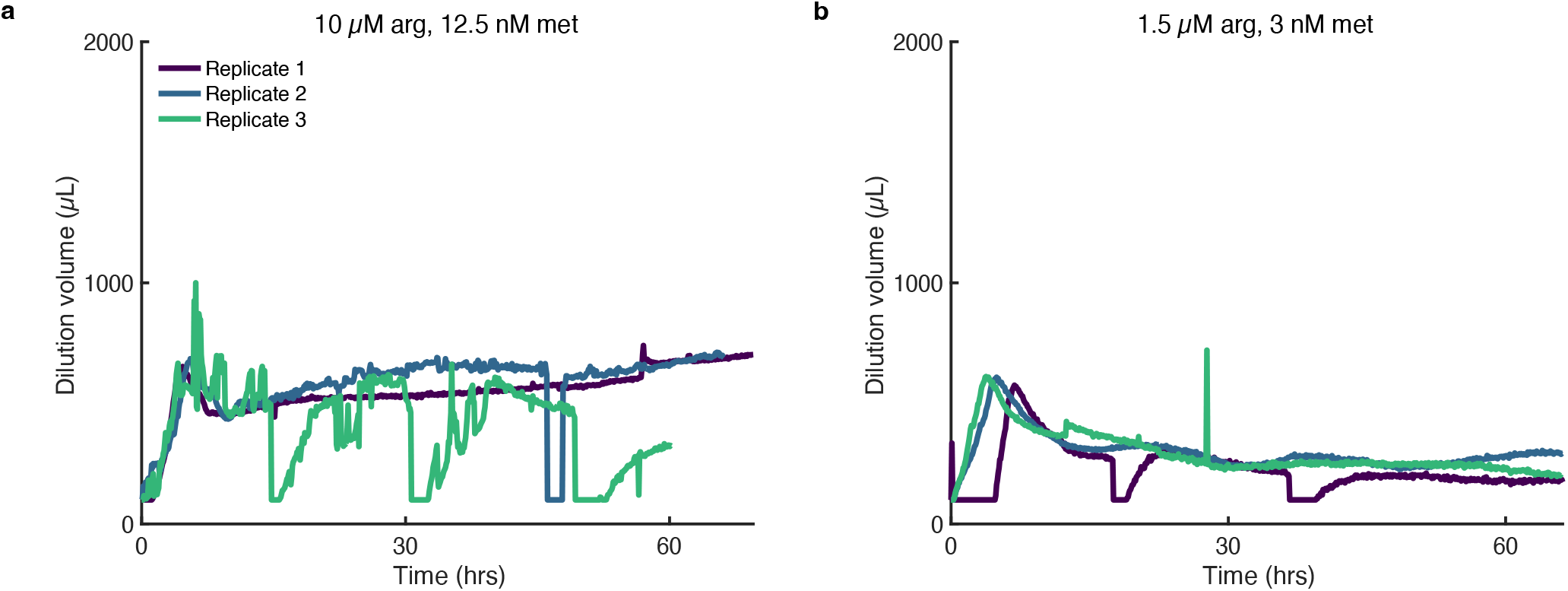
Turbidostat dilution volumes during prediction experiments. Dilutions calculated as in Supplementary Figure 5. Dilution plots correspond to ratio plots in Figure 4b-c. Spikes are due to misreads from OD chamber.

#### Phase 2

We fixed all the model parameters in the nonautonomous model, except *a* and *g*, to the parameter values found in Phase 1. We then fit *a* and *g* using the measurements taken over the first 15 hours in each of the six experiments. The priors for *a* and *g* are recorded in Table 2. The prior for *a* reflects the assumption that the initial constant growth regime is not weighted heavily after the first 15 hours of the experiment. The prior for *g* is set to assume that in the initial constant growth phase, the cells most likely grow more quickly than later in the experiment, but no more than twice as quickly. The posterior distribution is shown in Figure 7. We see that *a* and *g* are correlated, and we use the mean of the posterior (point indicated in the figure) to produce the figures in the main body of the text. Although we cannot resolve both *a* and *g* independently from the data, this model 1) provides evidence that the cells are in a different growth regime at the beginning of the experiments and 2) explains the steady states for every supplementation experiment. Importantly, the model also accurately predicts steady state values for the two new experiments shown in Figure 4.

### Observations about the posterior distribution

This section refers to the posterior of Phase 1 of the nonautonomous model fitting procedure, Figure 6.

The mode of the posterior distribution is 0 for the parameters *η*_*a*_ and *η*_*m*_, but nonzero values are also accepted in the posterior. Our intuition for this is the following: *η*_*a*_ (respectively *η*_*m*_) represents the degree to which arginine (resp. methionine) is consumed by Δ*ArgC* (resp. Δ*MetA*) cells. We expect *η*_*a*_ and *η*_*m*_ to be low compared to the production rates of each metabolite, especially because the dilution term in the model provides a second means by which each metabolite is removed from the system. Additionally, we find that the ratio *c*_*m*_(*t*) is relatively insensitive to choice in *η*_*a*_ and *η*_*m*_, as long as *η*_*a*_ and *η*_*m*_ are chosen to be in the interval indicated by the posterior distribution. We interpret this result to mean that, whereas the consumption rates of the metabolites are not necessarily 0, this mechanism contributes less to the resulting cell population proportion than other factors such as cell growth rate and metabolite production.

For the growth rate Hill coefficient *b*_*a*_, the mode of the posterior is near the highest endpoint of the prior. We have found that increasing *b*_*a*_ further does not influence the population ratio *c*_*m*_. Thus we use the mean value for *b*_*a*_ in our results, noting that other larger values for *b*_*a*_ will give similar results.

## Dataset_S1.xlsx

Colony counts of LB and M9-methionine plates for continuous culture experiments. Each sheet corresponds to one biological replicate.

## Dataset_S2.xlsx

Turbidostat recorded data for continuous culture experiments, including OD measurements and dilution volumes. Each sheet corresponds to one biological replicate.

## Dataset_S3.xlsx

Bulk culture OD timeseries data for monoculture metabolite growth response, displayed in Figure 1 C and D. Each column corresponds to one well of a 96-well plate, which contained either auxotroph and its corresponding metabolite at the labeled concentration.

